# Genetic and physical interactions between the organellar mechanosensitive ion channel homologs MSL1, MSL2, and MSL3 reveal a role for inter-organellar communication in plant development

**DOI:** 10.1101/487694

**Authors:** Josephine S. Lee, Margaret E. Wilson, Ryan A. Richardson, Elizabeth S. Haswell

## Abstract

Plant development requires communication on many levels, including between cells and between organelles within a cell. For example, mitochondria and plastids have been proposed to be sensors of environmental stress and to coordinate their responses. Here we present evidence for communication between mitochondria and chloroplasts during leaf and root development, based on genetic and physical interactions between three <u>M</u>echanosensitive channels of <u>S</u>mall conductance-<u>L</u>ike (MSL) proteins from *Arabidopsis thaliana*. MSL proteins are *Arabidopsis* homologs of the bacterial <u>M</u>echano<u>s</u>ensitive <u>c</u>hannel of <u>S</u>mall conductance (MscS), which relieves cellular osmotic pressure to protect against lysis during hypoosmotic shock. MSL1 localizes to the inner mitochondrial membrane, while MSL2 and MSL3 localize to the inner plastid membrane and are required to maintain plastid osmotic homeostasis during normal growth and development. In this study, we characterized the phenotypic effect of a genetic lesion in *MSL1*, both in wild type and in *msl2 msl3* mutant backgrounds. *msl1* single mutants appear wild type for all phenotypes examined. The characteristic leaf rumpling in *msl2 msl3* double mutants was exacerbated in the *msl1 msl2 msl3* triple mutant. However, the introduction of the *msl1* lesion into the *msl2 msl3* mutant background suppressed other *msl2 msl3* mutant phenotypes, including ectopic callus formation, accumulation of superoxide and hydrogen peroxide in the shoot apical meristem, decreased root length, and reduced number of lateral roots. All these phenotypes could be recovered by molecular complementation with a transgene containing a wild type version of *MSL1*. In yeast-based interaction studies, MSL1 interacted with itself, but not with MSL2 or MSL3. These results establish that the abnormalities observed in *msl2 msl3* double mutants is partially dependent on the presence of functional MSL1 and suggest a possible role for communication between plastid and mitochondria in seedling development.

## INTRODUCTION

Plastids and mitochondria are found in almost every plant cell and are involved in all aspects of plant biology. In plants, as in animals, mitochondria are involved in multiple cellular processes, including cellular respiration and co-enzyme synthesis (Schertl and Braun, 2014; Rébeillé et al., 2007). Plastids are responsible for photosynthesis and a range of other biosynthetic reactions—including the production of starch, some amino acids, fatty acids and lipids, pigments, hormones and volatiles (Neuhaus:2000eh; Rolland et al., 2018). Some plastids play a unique role in plant biology: amyloplasts in the root tip and the shoot endodermis are essential for gravity response (Toyota:2013fe; Su et al., 2017). A recent report argues that plastids of the leaf epidermis can serve as stress sensors (Beltrán et al., 2018). While individual reactions that take place in the plastid or mitochondrion benefit from their compartmentalization, broad metabolic processes are coordinated between them and the rest of the cell (Sweetlove and Fernie, 2013; Rolland et al., 2012; Schrader and Yoon, 2007). Furthermore, plastids and mitochondria physically interact with multiple other cellular compartments, including the nucleus, peroxisomes, and the ER (Kwok and Hanson, 2004; Barton et al., 2018; Kumar et al., 2018; Jaipargas et al., 2016; Mueller and Reski, 2015).

Metabolic integration between plastids and mitochondria is particularly intimate, especially under stress conditions (Raghavendra and Padmasree, 2003). For instance, the pool of cytoplasmic ATP is coordinately produced by chloroplasts and mitochondria; the extent to which each organelle contributes depends on current conditions (Gardeström and Igamberdiev, 2016). Mitochondria, chloroplasts, and peroxisomes collaborate extensively during photorespiration (Nunes-Nesi et al., 2008) (Hodges et al., 2016). Mitochondrial activity is thought to protect against photoinhibition and oxidative damage to chloroplasts by dissipating excess redox equivalents from the chloroplasts under high light conditions (Yoshida et al., 2007). Conversely, mitochondrial respiration has long been understood to be modulated by light. For example, the alternative oxidase AOX1a (a component of the mitochondrial electron transport chain) is up-regulated by light (Yoshida:2011ey; Yoshida et al., 2008).

The mechanism by which chloroplasts and mitochondria communicate is not fully understood. While there is evidence for the transfer of lipids via physical contact between chloroplasts and mitochondria (Jouhet et al., 2004), further validation is required (Delfosse et al., 2015). Communication may be mediated through the diffusion of factors through the cytosol, through direct contacts with other organelles (de Souza et al., 2017), or via signals to the nuclear genome (retrograde signaling) that are then conveyed to the other organelle (Kleine and Leister, 2016; de Souza et al., 2017; Chan et al., 2016; Woodson and Chory, 2012).

We have been studying the effect of organellar osmotic stress on plant development. We previously showed that two members of the MscS-Like (MSL) family of mechanosensitive ion channels, MSL2 and MSL3, serve to maintain osmotic homeostasis in plastids during normal growth and development (Veley et al., 2012; Haswell and Meyerowitz, 2006). MSL proteins are homologs of the mechanosensitive channel MscS (Mechanosensitive channel of small conductance), which serves as an “osmotic safety valve” to protect *Escherichia coli* against lysis during extreme hypoosmotic shock (Levina et al., 1999; Naismith and Booth, 2012). MSL2 and MSL3 are localized to the inner chloroplast membrane and *msl2 msl3* double mutants produce a range of plastid defects, including enlarged and round epidermal cell plastids, defective chloroplast division and abnormal ultrastructure in the proplastids of the shoot apical meristem (Wilson:2011dd; Wilson et al., 2014; 2016; Haswell and Meyerowitz, 2006). Furthermore, *msl2 msl3* plants have multiple developmental defects, including dwarfing and leaf variegation. After culture on solid media, they form ectopic calluses at the meristem, a process that is dependent on superoxide accumulation in plastids (Wilson et al., 2016). All of these developmental phenotypes can be interpreted as direct or indirect consequences of plastid osmotic dysregulation, as all are suppressed when plants or cells are supplied with osmotic support (Veley et al., 2014; Wilson et al., 2014; 2016). MSL2 and MSL3 can partially rescue a MS channel mutant *E. coli* strain, suggesting that they form MS ion channels as shown for several other members of the family (Haswell and Meyerowitz, 2006; Maksaev and Haswell, 2012; Lee et al., 2016),(Hamilton et al., 2015), but their electrophysiological characterization remains elusive.

MSL2 and MSL3 are two members of a ten-gene family in the genome of *Arabidopsis thaliana* (Haswell, 2007). Another member, MSL1, is also found in endosymbiotic organelles. Subcellular fractionation and GFP-fusion protein localization experiments demonstrate that MSL1 localizes to the inner mitochondrial membranes (Lee et al., 2016; Haswell and Meyerowitz, 2006). The mature form of MSL1 provides a mechanically activated ion channel activity in excised membrane patches (Lee et al., 2016). Plants harboring the null *msl1-1* allele (hereafter referred to as *msl1*) are indistinguishable from the wild type under normal growth conditions. However, plant mitochondria isolated from *msl1* mutants exhibit increased transmembrane potentials when the F_1_F_0_ATP synthase is inhibited. Compared to the wild type, *msl1* mutants also show a larger increase in mitochondrial glutathione oxidation in response to oligomycin, high temperature, and cadmium treatments, as measured with a redox-sensitive fluorescent reporter (mito-roGFP2). These data show that MSL1 plays a role in maintaining mitochondrial redox homeostasis during abiotic stress, but how direct these effects are and the role (if any) played by membrane stretch or ion flux is not yet known.

The presence of MSL channels in both chloroplast and mitochondrial envelopes, combined with existing evidence for integration of organellar responses to environmental and metabolic signals, led us to propose that MSL1, MSL2, and MSL3 may interact to coordinate a cellular response to osmotic stresses. To begin to test this idea, we characterized the genetic and physical interactions between MSL1, MSL2, and MSL3 in Arabidopsis. Our results reveal an unexpected genetic relationship whereby loss of *MSL1* enhances some but suppresses other phenotypes previously observed in the *msl2 msl3* mutant. We also document new phenotypes in the *msl2 msl3* root and show that these are also ameliorated in the *msl1 msl2 msl3* triple mutant. Finally, we demonstrate that MSL1 and MSL2 are capable of interacting with themselves in the split-ubiquitin yeast two hybrid assay, and that MSL2 and MSL3 interact with each other, but not with MSL1. These results point to a complex interplay between osmotic stress signals from the chloroplast and the mitochondria that lead to developmental outcomes in both the shoot and the root.

## METHODS

### Topology prediction and multiple sequence alignment

Sequences of *Ec*MscS, *At*MSL1 (Uniprot Q8VZL4), *At*MSL2 (isoform 1, Uniprot Q56X46) and *At*MSL3 (Uniprot Q8L7W1) were obtained from Uniprot (The UniProt Consortium, 2017). Mature MSL1 was defined as the protein remaining after cleavage of the mitochondrial transit peptide at Phe-79 (RAF↓SS; (Lee et al., 2016)), while mature MSL2 and MSL3 were defined as the protein remaining after cleavage of the predicted chloroplast transit peptide at Arg-75 (AFR↓CH) and Arg-70 (SSR↓CN) respectively (Haswell and Meyerowitz, 2006). Transmembrane domains and overall topology were predicted with Aramemnon (Schwacke and Flügge, 2018). Amino acid sequences were aligned using Clustal Omega 1.2.4 and default settings (Sievers and Higgins, 2018). Percent identity and similarity were calculated as number of identical or similar residues in the alignment divided by the total number of positions in the alignment, including gaps.

### Generation and validation of *msl1 msl2 msl3* triple mutant and *msl1 msl2 msl3 + MSL1g* complementation lines

The *msl1 msl2 msl3* triple mutant was generated by crossing the *msl1-1* mutant (first reported in (Lee et al., 2016)) to the *msl2-3 msl3-1* double mutant (first reported in (Wilson et al., 2011)). Triple mutant plants were identified in the F3 generation by PCR genotyping. A genomic copy of the *MSL1* locus (including all sequence from 1207 bp upstream of the ATG to 208 bp downstream of the TAG, including introns) was cloned into the pBGW backbone to make the molecular complementation construct *MSL1g* (Lee et al., 2016). To generate homozygous *msl1 msl2 msl3* lines complemented with a genomic copy of *MSL1* (*msl1 msl2 msl3* + *MSL1g*), *MSL1g* was introduced into the *msl1 msl2 msl3* background via Agrobacterium-mediated floral dip (Clough and Bent, 1998). Siblings homozygous for the presence or absence of *MSL1g* were identified in the T3 as lines exhibiting 100% or 0% Basta-resistance, respectively. All lines were validated by PCR genotyping. The MSL1 locus and our approach to genotyping the genomic locus of *MSL1* in the presence of *MSL1g* are shown in **Figure S1**.

### Plant growth

Plants were grown on soil at 23°C under a 16 h light regime (∼150 mmol m^−2^ s^−1^). For plants grown on solid media, seeds were surface-sterilized, stratified at 4°C in the dark for 2 days and placed on 1x Murashige and Skoog medium (pH 5.7; Caisson Labs) with 0.8% agar (Caisson Labs). They were grown vertically at 21°C under a 16-h-light regime with light fluence from 150 to 195 μmol m^−1^ s^−1^ for the indicated times.

### Superoxide and hydrogen peroxide detection

For superoxide detection, 21-day-old seedlings were collected, vacuum-infiltrated for 4 min in 0.1% weight-to-volume nitro blue tetrazolium in 10 mM K_2_HPO_4_-KH_2_PO_4_ potassium buffer pH 7.8 with 10 mM NaN_3_, incubated for 1 h in the dark, and then cleared with an ascending series of ethanol solutions (30%, 50%, 70%, 80% and 95%). This protocol was adapted from (Hoffmann et al., 2005). Images of stained seedlings were captured with a dissecting microscope and camera. Hydrogen peroxide detection was performed as described in (Wu et al., 2012) with the following modifications: seedlings were collected, incubated for 3 h in 0.1 mg/ml 3,3-diaminobenzidine pH 3.8, and vacuum-infiltrated for 5 min. Tissue was incubated overnight in the dark and cleared with an ascending ethanol series (30%, 50%, 70%, 80% and 95%), then imaged as for superoxide staining above.

### Mating-Based Split-Ubiquitin System

Physical interactions between MSL1, MSL2, and MSL3 were determined using the mating-based split-ubiquitin system described in (Obrdlik et al., 2004). *cDNA*s encoding the mature version of each protein were cloned and recombined into the destination vector *pEarleyGate103* (Earley et al., 2006) using LR Clonase II (Thermo Fisher Scientific). *MSL* sequences were PCR-amplified from destination vectors using primers attB1-F (5’-ACAAGTTTGTACAAAAAAGCAGGCTCTCCAACCACCATG-3’) and attB2-R (5’-TCCGCCACCACCAACCACTTTGTACAAGAAAGCTGGGTA-3’). PCR products were co-transformed with digested *pMetYCgate* (digested with *EcoRI+SmaI*) into yeast strain THY.AP4 (selected on Synthetic Complete media lacking leucine), and with digested *pXNGate21-3HA* (digested with *PstI+HindIII*) into yeast strain THY.AP5 (selected on Synthetic Complete media lacking tryptophan and uracil). *pMetYCgate* and *pXNGate21-3HA* were obtained from the Arabidopsis Biological Resource Center. Cells were mated for two days on Synthetic Complete media lacking Leu, Trp, and Ura for selection of diploids. Interactions between proteins were determined via growth after three days on Synthetic Minimal media lacking adenine, histidine, leucine, tryptophan, and uracil, and supplemented with 150 μM Methionine.

## RESULTS

### Topological comparison of *Escherichia coli* MscS and organellar *Arabidopsis thaliana* MscS-Like monomers

Both crystallography and biochemical experiments establish that *Ec*MscS forms a homoheptameric mechanosensitive ion channel (Bass:2002hg; Miller et al., 2003)}. Each *Ec*MscS monomer contributes three transmembrane (TM) domains and a relatively large soluble cytoplasmic domain. Like other MscS-like superfamily proteins, MSL1, MSL2 and MSL3 share a conserved region corresponding to the pore-lining helix and about 100 amino acids of the cytoplasmic C-terminus called the MscS domain ((Basu and Haswell, 2017), indicated in yellow in **Figure 1**). Outside of this domain, the topology of organellar MSL channels differ from *Ec*MscS and from each other in a number of ways. MSL1, 2, and 3 all are larger than MscS and have five TM domains with internal and external loops. MSL1 has an extended soluble N-terminal domain, while MSL2/3 have an extended C-terminal domain (only MSL2 is shown in **Figure 1**). Mitochondrial fractionation experiments suggest that the preprotein version of MSL1 is targeted to mitochondria by the N-terminal targeting peptide (Lee et al., 2016) (indicated in red), which is proteolytically cleaved after organellar import. Similarly, it is likely that the chloroplast-targeting N-terminal peptides of MSL2 and MSL3 (indicated in green) are cleaved after directing preprotein to the chloroplast (Haswell and Meyerowitz, 2006).

**Figure 1.**
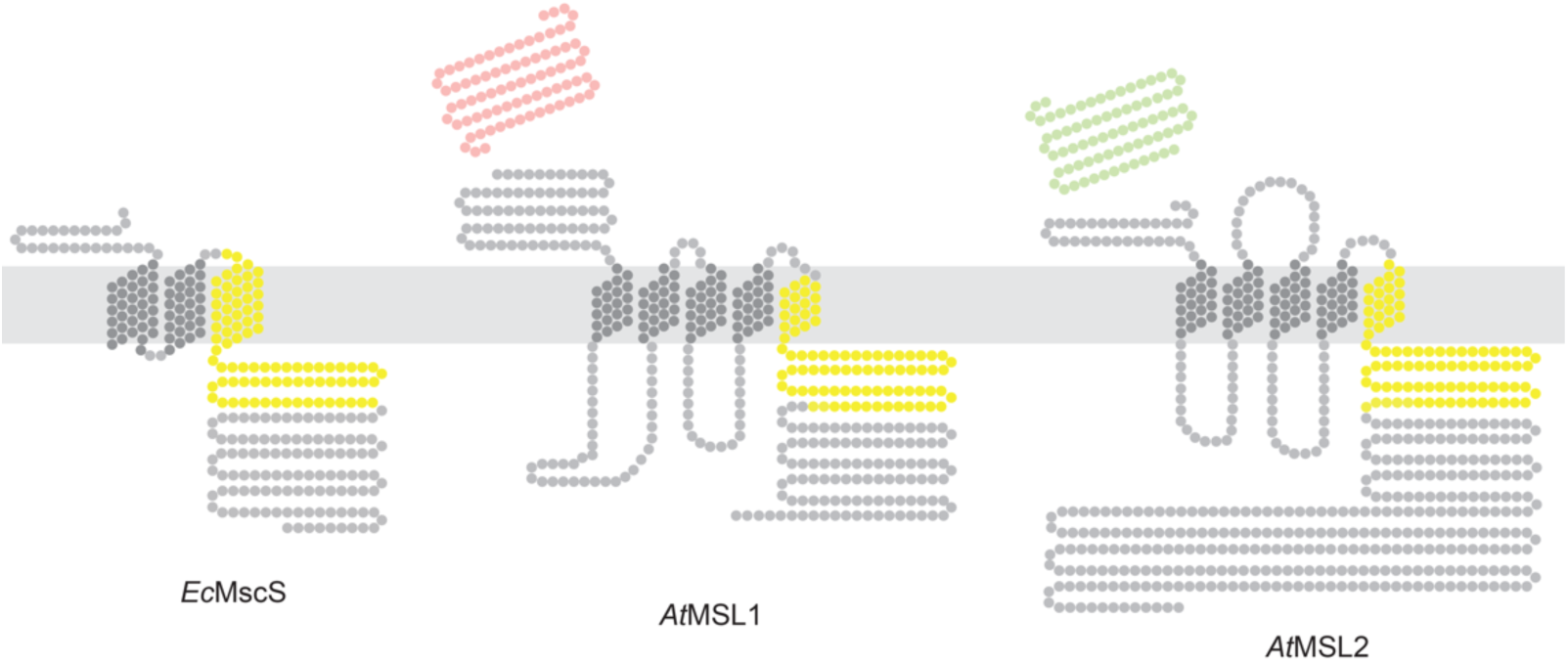
Predicted Topology of *Ec*MscS, *At*MSL1, and *At*MSL2. Experimentally determined or predicted membrane topology of the indicated monomers. Each dot represents one amino acid. Amino acids corresponding to the conserved MscS Domain (as defined in (Haswell, 2007)) are indicated in yellow; the MSL1 mitochondria targeting peptide (as defined in (Lee et al., 2016)) is indicated in red; and the MSL2 chloroplast targeting peptide (as defined in (Haswell and Meyerowitz, 2006)) is shown in green.

### Loss of *MSL1* exacerbates the leaf notching, rumpling and variegation observed in *msl2 msl3* double mutant plants

In order to address the possibility of coordination between plastids and mitochondria, we first investigated genetic interactions between *MSL1, MSL2*, and *MSL3*. To do so, we compared the whole seedling phenotypes of 24-day-old wild type plants, *msl1* mutants, *msl2 msl3* double mutants, *msl1 msl2 msl3* triple mutants, and *msl1 msl2 msl3* triple mutants complemented with a transgene containing a genomic copy of *MSL1* (*msl1 msl2 msl3 + MSL1g*) (**Figure 2**). As previously reported, *msl2 msl3* plants exhibit leaf notching, rumpling and variegation (Wilson et al., 2011). While plants lacking functional *MSL1* appeared wild type, *msl1 msl2 msl3* triple mutant seedlings showed exacerbated leaf notching, rumpling and variegation compared to *msl2 msl3* double mutant seedlings. This effect was suppressed in *msl1 msl2 msl3* + *MSL1g* seedlings, indicating that the increase in phenotypic severity in the *msl1 msl2 msl3* triple mutant can be attributed to a defect at the *MSL1* locus.

**Figure 2.**
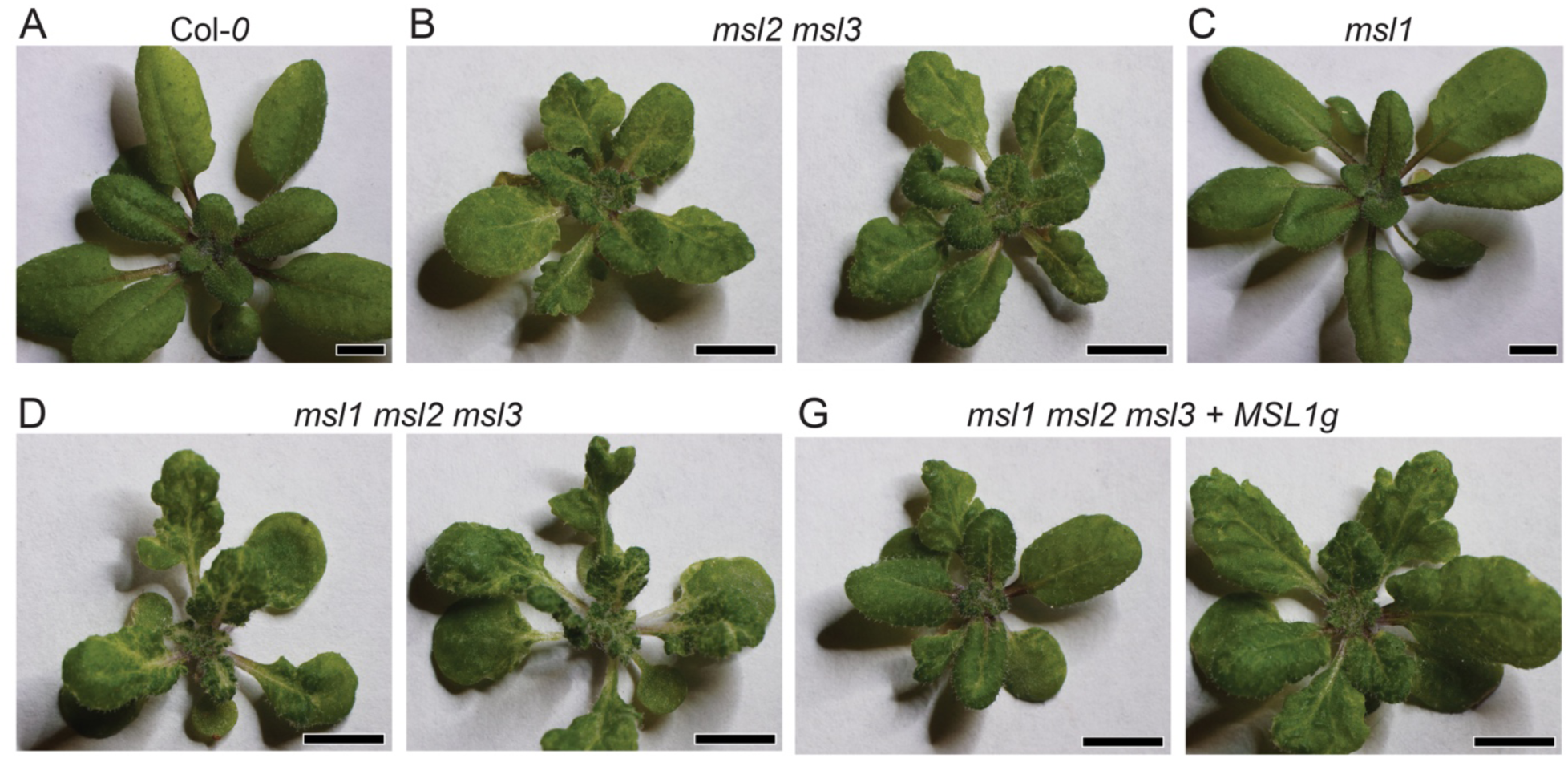
Loss of *MSL1* exacerbates the leaf notching, rumpling and variegation observed in *msl2 msl3* double mutant plants. Images of 24-day-old soil-grown seedlings of the following genotypes: (A) Col*-0*, (B) *msl2 msl3*, (C) *msl1* (D) *msl1 msl2 msl3*, and (E) *msl1 msl2 msl3 + MSL1g* plants. The scale bar represents 0.5 cm.

### *msl1 msl2 msl3* triple mutants form shooty outgrowths in place of the ectopic calluses observed in *msl2 msl3* double mutants

Since the *msl1* lesion exacerbated leaf phenotypes in the *msl2 msl3* background, we hypothesized that the same would be true for other *msl2 msl3* phenotypes, including the production of meristematic callus previously observed in *msl2 msl3* seedlings grown on solid media (Wilson et al., 2016). Seedlings were grown vertically on solid media for 19-21 days at 21°C under a 16-hour-light regime and the shoot apex examined **(Figure 3)**. Under these conditions, *msl1* seedlings were indistinguishable from the wild-type and meristematic calluses were not observed in either background. Consistent with our earlier report, callus-like growth at the shoot apex was observed in ∼70% of *msl2 msl3* seedlings. Unexpectedly, no callus was formed in over 180 *msl1 msl2 msl3* triple mutant plants examined. Instead, shooty outgrowths at the meristem were observed in 40-60% of these seedlings. Shooty outgrowths were never observed in *msl2 msl3* plants, nor in *msl1 msl2 msl3* + *MSL1g* plants, and the production of callus was recovered in *msl1 msl2 msl3* + *MSL1g* seedlings (88 of 131). Thus, *MSL1* is required for the formation of callus in *msl2 msl3* mutants, and in its absence, shoot-like growths are formed.

**Figure 3.**
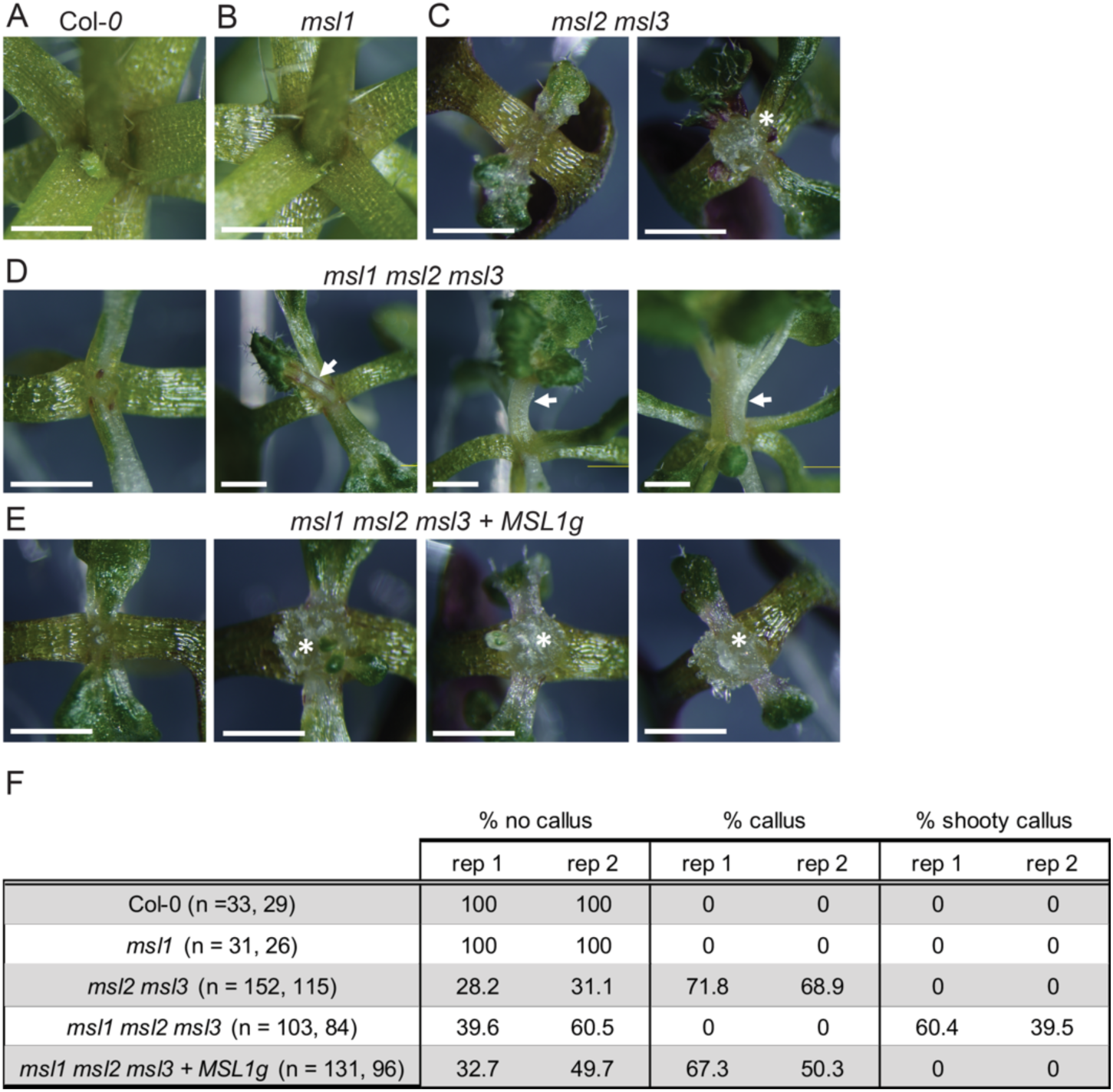
Addition of the *msl1* lesion to the *msl2 msl3* background causes formation of shooty outgrowths in place of ectopic calluses. Close-up images of the shoot apex of seedlings grown vertically on 1x MS media for 21 days. (A) Col-*0*, (B) *msl1*, (C) *msl2 msl3* with no callus (left) and callus (right); (D) *msl1 msl2 msl3* with no shooty outgrowth (left) and shooty outgrowths (center and right); and (E) *msl1 msl2 msl3 + MSL1g* with no callus (left) and callus (center and right). Asterisks indicate callus; arrows indicate shooty outgrowths. The scale bar represents 1 mm. (F) Percentage of seedlings exhibiting no callus, callus, and shooty callus in the indicated genotypes. Results from two independent experiments are shown and the number of seedlings included in each is indicated.

### *MSL1* is required for meristematic reactive oxygen species accumulation in the *msl2 msl3* background

Double *msl2 msl3* mutants accumulate the reactive oxygen species (ROS) superoxide (O_2_^−^) and hydrogen peroxide (H_2_O_2_) at the shoot apex at levels higher than the wild type (Wilson et al., 2016). To determine the role of *MSL1* in the accumulation of ROS, seedlings were grown on solid media for 21 days and stained with 3,3’-diaminobenzidine (DAB, which indicates H_2_O_2_) or nitrotetrazolium blue chloride (NBT, which indicates O_2_^−^) (**Figure 4**). As previously observed, levels of NBT and DAB were higher in *msl2 msl3* mutants shoot apices than in the wild type. Single *msl1* mutants were indistinguishable from the wild type. In the apices of *msl1 msl2 msl3* triple mutants, DAB and NBT staining were greatly reduced compared to *msl2 msl3* double mutants. In addition, strong meristematic DAB and NBT staining was recovered in *msl1 msl2 msl3 + MSL1g* plants, indicating that *MSL1* is required for meristematic ROS accumulation in addition to callus formation in *msl2 msl3* plants. These results are also consistent with our previous observation that callus formation in *msl2 msl3* seedlings is dependent on O_2_^−^ accumulation in the shoot apex (Wilson et al., 2016), and we propose that the lack of callus formation in *msl1 msl2 msl3* plants can be attributed to the absence of apical O_2_^−^ accumulation when *MSL1* is mutated. We note that NBT (but not DAB) staining in the cotyledons and leaves of the *msl1 msl2 msl3* triple mutant was elevated compared to all other genotypes (**Figure S2**).

**Figure 4.**
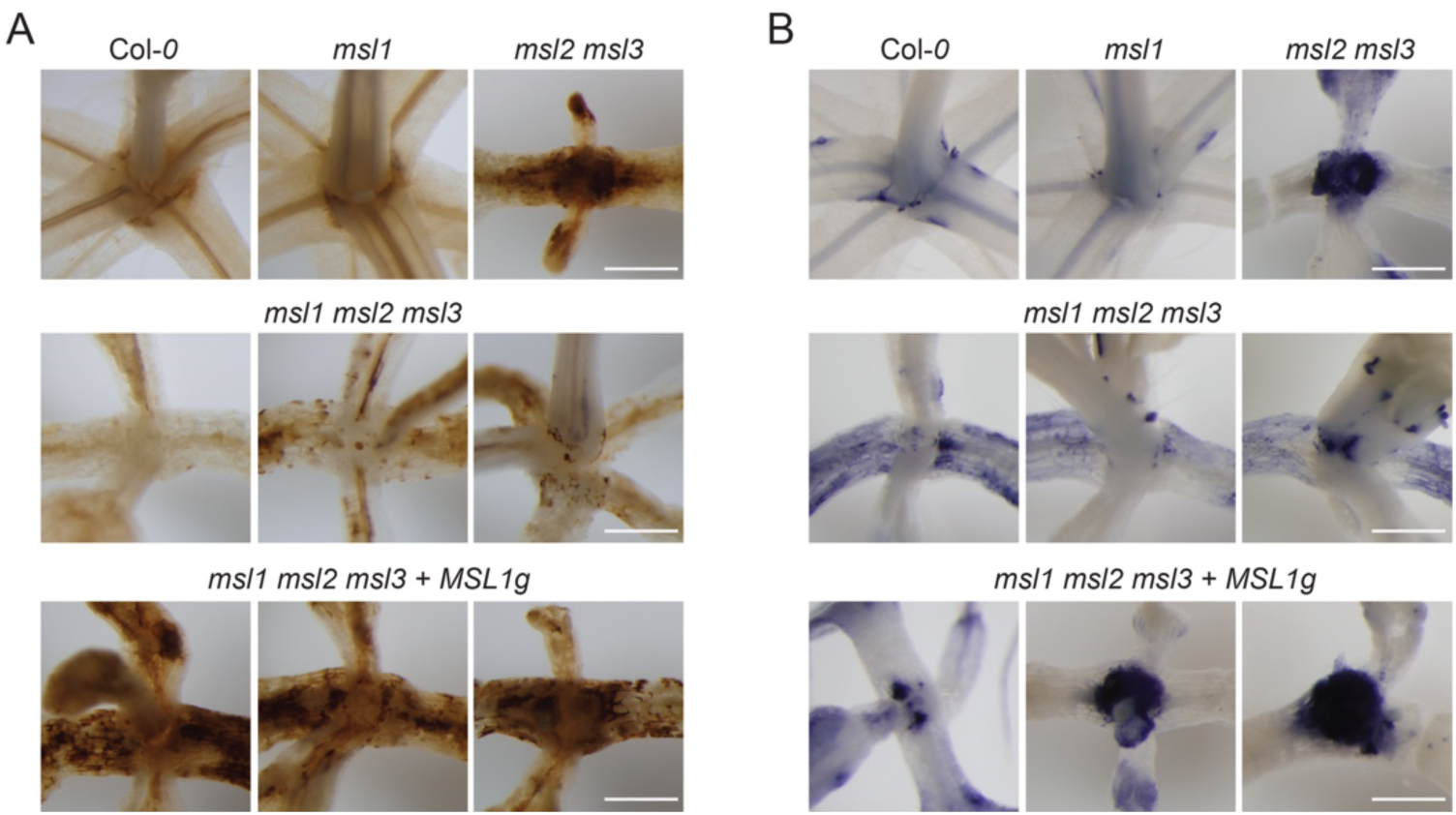
The *msl1* lesion suppresses meristematic ROS accumulation in the *msl2 msl3* background. Images of seedlings infiltrated with (A) 3,3’-diaminobenzidine (DAB) stain to visualize H_2_O_2_ accumulation and (B) nitrotetrazolium blue chloride (NBT) stain to visualize O^2-^ accumulation, then cleared in ethanol. All seedlings were grown vertically on 1X MS media for 21 days. The scale bars represent 0.5 mm.

### *msl2 msl3* mutants have shorter roots and few lateral roots per unit length, and *MSL1* is partially required for these root defects

Only aerial phenotypes of the *msl2 msl3* mutant have been documented. To begin to assess root phenotypes in this mutant, we grew seedlings vertically on solid media for 13 days. As shown in **Figure 5**, *msl2 msl3* seedlings had primary roots averaging 1.4 cm in length, over 4 times shorter than Col-*0* roots, which averaged 6.8 cm. Additionally, *msl2 msl3* mutants formed very few lateral roots, averaging 0.39 lateral roots/cm compared to the wild type average of 2.4 lateral roots/cm. We further observed that *msl1* mutant roots were 6.3 cm long and had 2 lateral roots/cm on average, comparable to that of wild type. *msl1 msl2 msl3* mutant roots were significantly longer than those of *msl2 msl3* seedlings with an average root length of 2.6 cm. They also had an average of 2.4 lateral roots/cm, statistically grouping with the wild type and significantly different from the average for *msl2 msl3* seedlings. *msl1 msl2 msl3 + MSL1g* seedlings had shorter root lengths averaging 1.3 cm that statistically grouped with those of *msl2 msl3* seedlings. They had an average of 1.3 lateral roots/cm, intermediate between that of *msl2 msl3* and wild type, and in a statistically separate group. In summary, the primary roots of *msl2 msl3* seedlings are shorter than the wild type with fewer lateral roots per cm. Further, *MSL1* is required for the observed short root phenotype, and appears to be involved in the reduction in lateral roots.

**Figure 5.**
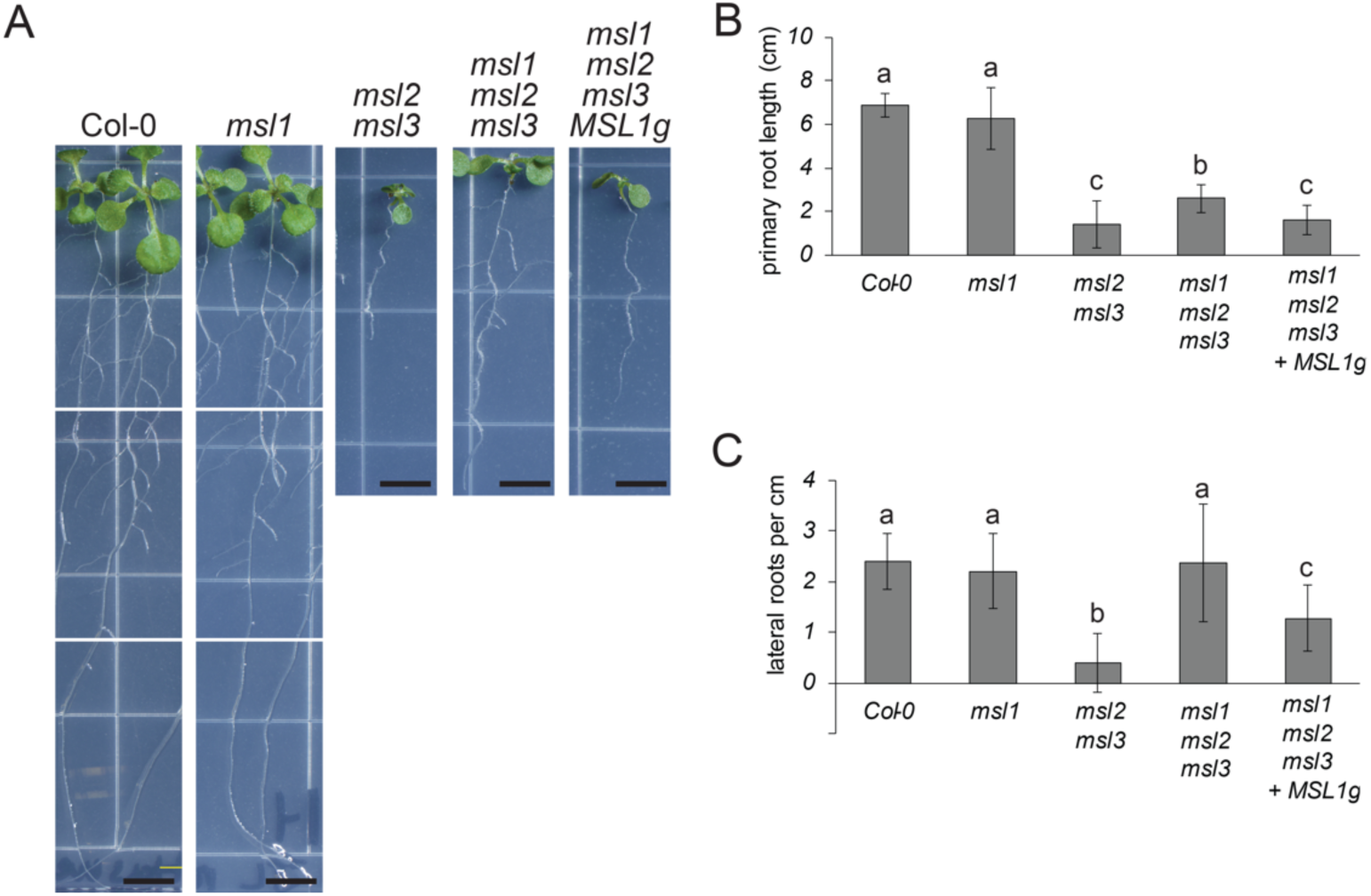
*msl2 msl3* mutants have shorter roots and fewer lateral roots than the wild type. These root defects are suppressed in the *msl1 msl2 msl3* background. (A) Representative images of seedlings grown vertically on 1x MS media for 13 days. The scale bar represents 0.5 cm. Quantification of (B) root length and (C) number of lateral roots per cm in seedlings grown as in (A) Error bars represent standard deviation. N = 24-36 seedlings per genotype. One-way analysis of variance (ANOVA) was used with a *p* < 0.05 cutoff for significance. Scheffe’s test was then used for post-hoc means separation, again with a *p* < 0.05 cutoff. Letters indicate different statistical groups using Scheffe’s test. Similar results were obtained in an independent experiment.

### MSL1, MSL2, and MSL3 interact in an organelle-specific manner

To begin to assess whether these genetic relationships might be mediated through direct protein-protein interactions, we used the mating-based split-ubiquitin system (mbSUS), a version of the classic yeast two hybrid modified for the analysis of membrane protein-protein interactions (Grefen et al., 2009; Obrdlik et al., 2004). In this assay, interactions between proteins are assessed by virtue of their ability to bring together two fragments of ubiquitin, Nub and Cub. When the two fragments are brought together, they catalyze the cleavage of an artificial transcription factor (LexA-VP16) that is translationally fused to Cub, thereby allowing activation of reporter genes. We tested mature (lacking transit peptides) versions of MSL1, MSL2, and MSL3 for interaction in this assay (**Figure 6)**. Mating yeast strains expressing MSL1, 2, or 3-Cub-LexA to a strain expressing Nub^WT^, a version of Nub that does not require interaction for growth, led to growth on drop-out media. Mating them to a strain with an empty NubG vector did not. We observed that MSL1-Cub-LexA interacted with MSL1-NubG, but not with MSL2-NubG nor MSL3-NubG. On the other hand, MSL2-Cub-LexA interacted strongly with MSL2-NubG and MSL3-NubG. MSL3-Cub-LexA only interacted with MSL2-NubG. In summary, MSL1 and MSL2 interacted with themselves, as expected for the monomers of multimeric channels. MSL2 and MSL3 also interacted with each other, implying the formation of heteromeric channels in the chloroplast envelope. However, MSL1 did not interact with MSL2 or with MSL3, and MSL3 did not interact with itself.

**Figure 6.**
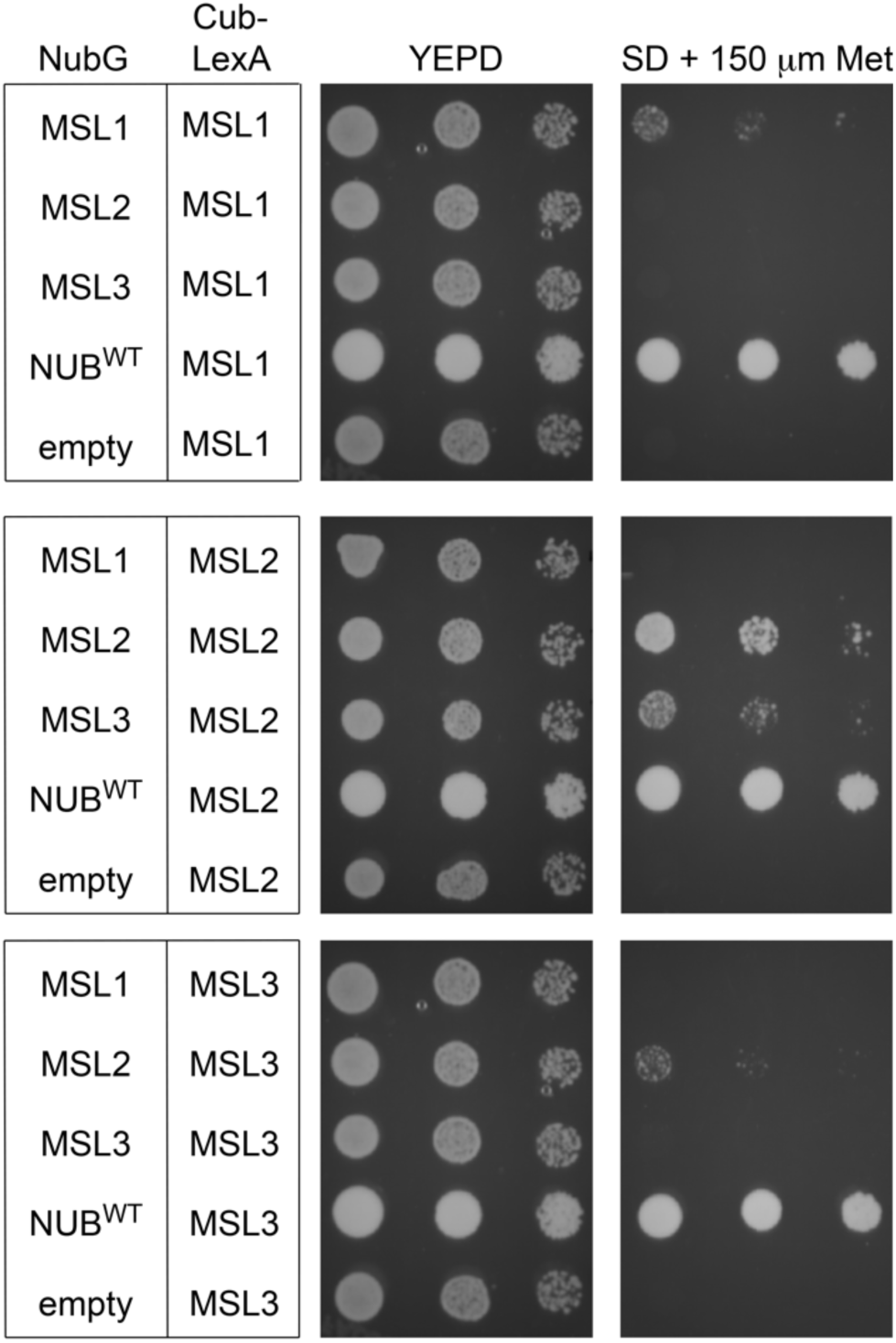
Mating-based split ubiquitin assay. Growth of diploids transformed with the constructs indicated on the left on YEPD or Synthetic Minimal media + 150μM Methionine. Left to right for each plate is one tenfold dilution (OD_600_ = 1.0, 0.1, 0.01). Growth assays were independently repeated 3 times.

## DISCUSSION

It has been proposed that plastids and mitochondria interact through signaling or metabolic pathways to coordinate cellular responses (Bobik and Burch-Smith, 2015), prompting us to initiate an analysis of the genetic and physical interactions between three members of the MscS-Like (MSL) family of mechanosensitive ion channels. These three proteins are localized to the mitochondria (MSL1 (Lee et al., 2016)) or to the chloroplast (MSL2 and MSL3 (Haswell and Meyerowitz, 2006)). While *msl1* mutant plants have no obvious developmental phenotype, *msl2 msl3* mutants exhibit crumpled, variegated, and notched leaves and after growth on solid media they produce callus at the shoot apex. Double *msl2 msl3* mutants also accumulate ROS at the shoot apex. Here we document two additional phenotypes in the *msl2 msl3* mutant, including a shorter primary root and reduced number of lateral roots than the wild type. In addition, we found that introducing the *msl1* allele into the *msl2 msl3* background exacerbated leaf phenotypes but ameliorated callus production, ROS accumulation, and the root phenotypes.

There are multiple molecular explanations for genetic interactions between proteins localized to different compartments. One possibility is that they are actually not in different compartments; that MSL1 could move to the chloroplast or MSL2 can move to the mitochondrion. Dual targeting to both the mitochondria and the chloroplast has been observed for many plant proteins but is difficult to predict (Carrie:2013kh; Xu et al., 2013). We have not observed dual localization in our experiments with MSL1-, MSL2-, or MSL3-GFP fusion proteins, but it remains a possibility that protein levels below the level of detection are dual localized. We considered the possibility of the formation of heteromeric channels, which might explain cross-organelle effects with very low levels of dual-targeted proteins. However, in our mbSUS experiments, we did not observe any interactions between MSL1 and MSL2 or MSL3, though we did see robust interaction between MSL2 and MSL2, and also strong interaction between MSL2 and MSL3 (**Figure 6**). Whether MSL3 forms a homomeric channel or is only able to form a heteromeric channel with MSL2 remains to be determined. Taken together, these data suggest that the observed genetic interactions between MSL1, MSL2, and MSL3 are unlikely to be mediated by direct protein-protein interactions.

Instead, the MSL1/2/3 genetic interactions we observed may reflect an interaction between two organelle signaling pathways that impinge on developmental outcomes such as leaf and root morphology and the differentiation of cells at the shoot apex. Double *msl2 msl3* mutant plastids are enlarged under osmotic stress. We’ve previously shown that the resulting phenotypes can be suppressed by growth on osmotica, establishing that they are produced in response to plastid osmotic stress. All aspects of the *msl2 msl3* phenotype: leaf morphology (**Figure 2**), ectopic callus (**Figure 3**), ROS accumulation (**Figure 4**), and short root and low number of lateral root phenotypes (**Figure 5**) were altered in the absence of *MSL1*, indicating that the signal or signals that induce these phenotypes require the presence of MSL1 and thus go through the mitochondria. Our current working hypothesis is that that *msl2 msl3* mutant plastids produce or potentiate an osmotic stress signal that requires MSL1 function in the mitochondria for its production or action. When MSL1 is absent, the osmotic stress signal generated in the plastids is not propagated, resulting in exacerbated (leaf morphology) or attenuated (callus, ROS, root) phenotypes.

With respect to callus production, one mechanism by which mitochondria might affect plastid osmotic stress signaling is through the modulation of ROS levels. We previously showed that ROS accumulation in the shoot apex leads to and is required for apical callus formation in *msl2 msl3* plant, (Wilson et al., 2016). Furthermore, *msl1* mutants show increased mito-roGFP signal in response to multiple abiotic stresses, indicating that the mitochondrial glutathione pool is oxidized under these conditions (Lee et al., 2016). We proposed that MSL1 is required to prevent over-reduction of the respiratory chain and ROS production under conditions of high membrane potential. The data presented here suggest that a mitochondrial signal associated with MSL1 functions upstream of meristematic superoxide accumulation and the production of callus that is caused by osmotically stressed plastids in the *msl2 msl3* mutant. One possibility is that osmotically stressed plastids in some way induce a ROS-related stress signal in mitochondria, which turn leads to the accumulation of ROS in meristematic cells and the production of callus. We propose that in the absence of MSL1, the signal from plastids is not efficiently received or propagated, perhaps because *msl1* mutant mitochondria are unable to normalize their own ROS levels and therefore have an abnormal response to a subtle ROS signal from osmotically stressed plastids.

To summarize, we show here that the loss of *MSL1* can attenuate or exacerbate the developmental effects of plastid osmotic stress observed in the *msl2 msl3* mutant. We hypothesize a signaling relationship between the two organelles that impacts a range of developmental processes, from cell identity at the shoot apex to the elaboration of lateral roots. Additional experiments are needed to determine how osmotically stressed plastids lead to these developmental phenotypes, and why many of them are modulated by the presence of mitochondrial MSL1.

---

**Figure S1. Primers involved in genotyping the genomic locus of *MSL1* in the presence of the *MSL1g* transgene.** (A) Schematic of the *MSL1* gene and the location of primers used. Thick lines indicate exons and thin lines indicate introns. The inverted triangle indicates the insertion point of the T-DNA in the *msl1-1* mutant. Thin arrows (not to scale) indicate the approximate recognition sites of oligos used for genotyping. (B) Primer pairs used to distinguish the wild type genomic version of *MSL1*, the *MSL1g* transgene, and the *msl1-1* T-DNA insertion allele. (C) The sequences of primers used in (B) are shown.

**Figure S2.** Images of seedlings infiltrated with (A) 3,3’-diaminobenzidine (DAB) stain to visualize H_2_O_2_ accumulation and (B) nitrotetrazolium blue chloride (NBT) stain to visualize O^2-^ accumulation, then cleared in ethanol. All seedlings were grown vertically on 1x MS media for 21 days. The scale bar represents 6 mm.

## ACKNOWLEDGEMENTS

This research was supported by an American Society of Plant Biologists Summer Undergraduate Research Fellowship to JSL and NSF MCB-1253103 to ESH. The research of RR was supported in part by a Faculty-Scholar grant to ESH from the Howard Hughes Medical Institute and the Simons Foundation. We thank the Jeanette Goldfarb Plant Growth Facility staff for assistance with plant growth and Angela Schlegel for help with supplementary figure preparation.

## AUTHOR CONTRIBUTIONS

MEW and ESH designed the research and supervised the experiments. JSL and RR performed the experiments and JSL and ESH drafted the manuscript with contributions from RR and MEW. MEW generated the *msl1 msl2 msl3* mutant and made initial observations, and RR conducted the mating-based split ubiquitin experiments. JSL, MEW RR, and ESH analyzed data.

